# Ion permeation pathway within the internal pore of P2X receptor channels

**DOI:** 10.1101/2022.11.22.517476

**Authors:** Stephanie W. Tam, Kate Huffer, Mufeng Li, Kenton J. Swartz

## Abstract

P2X receptor channels are trimeric ATP-activated ion channels expressed in neuronal and non-neuronal cells that are attractive therapeutic targets for human disorders. Seven subtypes of P2X receptor channels have been identified in mammals that can form both homomeric and heteromeric channels. P2X1-4 and P2X7 receptor channels are cation selective, whereas P2X5 has been reported to have both cation and anion permeability. P2X receptor channel structures reveal that each subunit is comprised of two transmembrane helices, with both N-and C-termini on the intracellular side of the membrane, and a large extracellular domain that contains the ATP binding sites at subunit interfaces. Recent structures of ATP-bound P2X receptors with the activation gate open reveal the unanticipated presence of a cytoplasmic cap over the central ion permeation pathway, leaving lateral fenestrations that may be largely buried within the membrane as potential pathways for ions to permeate the intracellular end of the pore. In the present study we identify a critical residue within the intracellular lateral fenestrations that is readily accessible to thiol reactive compounds from both sides of the membrane and where substitutions influence the relative permeability of the channel to cations and anions. Taken together, our results demonstrate that ions can enter or exit the internal pore through lateral fenestrations that play a critical role in determining the ion selectivity of P2X receptor channels. (223)

## INTRODUCTION

P2X receptors are a family of ion channels that are activated by extracellular ATP (Khakh and North 2006). There are seven subtypes of P2X receptor channels that are widely expressed throughout the body, including the central and peripheral nervous systems, as well as the cardiovascular, immune, respiratory, gastrointestinal, and genitourinary systems (Khakh and North 2006, Illes, Muller et al. 2021). They are thought to play a wide range of important roles, from transmitting gustatory signals, sensing bladder filling, mediating neuropathic and inflammatory pain, to regulation of immune responses (Khakh and North 2006, Surprenant and North 2009, Schmid and Evans 2019, Illes, Muller et al. 2021).

Structures of P2X3, P2X4 and P2X7 receptor subtypes solved using X-ray crystallography or cryo-electron microscopy (cryo-EM), reveal that they are trimers with each subunit containing two transmembrane (TM) helices, with both N- and C-termini on the intracellular side of the membrane and a large extracellular domain that contains the ATP binding sites at the subunit interfaces (Fig. 1A) (Kasuya, Fujiwara et al., Kawate, Michel et al. 2009, Hattori and Gouaux 2012, Mansoor, Lu et al. 2016, Li, Wang et al. 2019, McCarthy, Yoshioka et al. 2019). The TM2 helix lines the ion permeation pathway, with the activation gate positioned towards the external end of the pore (Li, Chang et al. 2008, Kawate, Michel et al. 2009, Li, Kawate et al. 2010, Hattori and Gouaux 2012). Most P2X receptors are permeable to cations like Na^+^ and Ca^2+^ (North 2002, Egan and Khakh 2004); however, the P2X5 receptor has been reported to have measurable Cl^-^ permeability (Ruppelt, Ma et al. 2001, Bo, Jiang et al. 2003). Although important determinants influencing the relative permeability of Na^+^ and Ca^2+^ have been identified within the external pore of P2X receptors (Migita, Haines et al. 2001, Samways and Egan 2007, Samways, Li et al. 2014), the regions of the pore determining the relative permeability of cations over anions are unknown.

**Figure 1.**
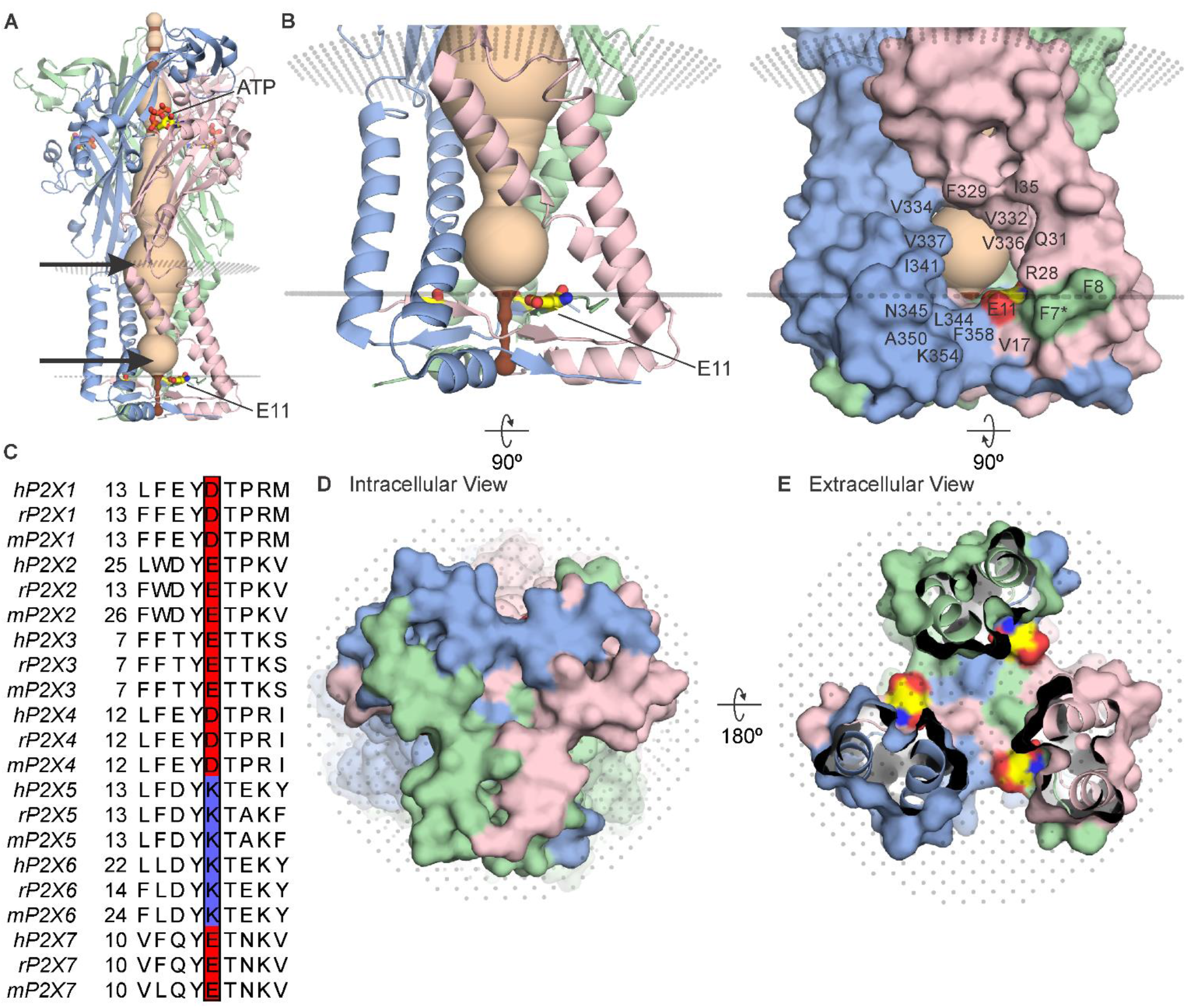
The cytoplasmic cap in P2X3 receptor channels (7 inch) **(A)** Side view of the structure of hP2X3_Slow_ with ATP bound (PDB ID: 6ah4). Ribbon representations of each subunit are colored blue, pink and green, with a HOLE representation along the central axis colored with radii ≤ 2 Å in brown and radii larger than 2 Å in tan. E11 (corresponding to E17 in rP2X2) is shown in stick representation with carbon (yellow), nitrogen (blue) and oxygen (red). OPM representation of the membrane is shown in light gray spheres and location of lateral fenestrations at both extracellular and intracellular ends of the pore indicated with arrows. **(B)** Magnified side view of the transmembrane helices and cytoplasmic cap shown in ribbon representation (left) and with surface representation (right). Residues around the lateral fenestrations are labeled. **(C)** Multiple sequence alignment of all human, rat, and mouse P2X subtypes for the region of the cap containing E17 in rP2X2. Residues corresponding to E17 in rP2X2 are outlined with a box, with acidic residues colored red and basic residues colored blue. A full sequence alignment for elements of the cytoplasmic cap is provided in Figure 2 – Figure Supplement 1 and Uniprot accession numbers are provided in the legend to Figure 2 – Figure Supplement 2. **(D)** Intracellular view of the cytoplasmic cap with surface representation. **(E)** View of the cytoplasmic cap from within the pore with surface representation and carbon colored yellow, nitrogen blue and oxygen red for atoms in E11.

All the available structures of P2X receptors contain lateral fenestrations between the TM and extracellular domains, providing a pathway for ions to permeate through the external end of the pore (Kawate, Robertson et al. 2011, Samways, Khakh et al. 2011)(Fig. 1A). Although the initial structures of P2X4 receptor channels contained an internal pore where ions could enter or exit at the three-fold axis of symmetry (Kawate, Michel et al. 2009, Hattori and Gouaux 2012), the N- and C-termini in these structures were not resolved. More recent structures of a slowly desensitizing mutant of the P2X3 receptor channel with ATP bound (Mansoor, Lu et al. 2016) and of the P2X7 receptor channel with and without ATP bound (Mansoor, Lu et al. 2016) reveal the presence of an intracellular structural element formed by two N-terminal ß-strands and a C-terminal α-helix and ß-strand that physically caps the cytoplasmic end of the pore (Fig. 1A, B, D, E). Although small lateral fenestrations present in the structures containing these cytoplasmic caps have been proposed to serve as the pathway for water and ions to enter or exit the internal pore (Mansoor, Lu et al. 2016), these lateral fenestrations would be expected to be largely buried within the membrane given the dimensions of the TM domains in P2X receptor channels (Mansoor, Lu et al. 2016, McCarthy, Yoshioka et al. 2019). However, the precise disposition of the lateral fenestrations relative to the membrane remains uncertain because these structures were solved in detergent solution (Mansoor, Lu et al. 2016, McCarthy, Yoshioka et al. 2019).

In the present study we explored whether the intracellular lateral fenestrations provide a pathway for ions to enter and exit the internal pore. We identified a residue lining the lateral fenestrations that is a conserved acidic residue in the cation selective P2X1-4 and P2X7 receptor channels but is a conserved basic residue in the anion permeable P2X5 receptor channel and could therefore play a role in determining ion selectivity. When mutated to Cys, this position is accessible to Cys-reactive reagents from both sides of the membrane, and charge-reversing mutations influence the relative permeability of cations to anions. Our results demonstrate that the lateral fenestrations provide an ion permeation pathway at the intracellular end of the pore and that they contain a critical determinant of ion selectivity.

## RESULTS

The objective of the present study was to investigate how ions permeate through the internal pore of P2X receptor channels given the presence of the cytoplasmic caps recently identified in P2X3 and P2X7 receptor channels (Mansoor, Lu et al. 2016, McCarthy, Yoshioka et al. 2019). Although the structures of P2X receptors solved thus far are remarkably similar within the large extracellular domain and relatively small TM domain (Kasuya, Fujiwara et al., Kawate, Michel et al. 2009, Hattori and Gouaux 2012, Mansoor, Lu et al. 2016, Li, Wang et al. 2019, McCarthy, Yoshioka et al. 2019), the cytoplasmic cap has only recently been resolved in P2X3 and P2X7 structures (Mansoor, Lu et al. 2016, McCarthy, Yoshioka et al. 2019)(Fig. 1; Fig. 2; Fig. 2 – Figure Supp. 1). The slowly desensitizing hP2X3 receptor structural construct (hP2X3_Slow_) where the cap was resolved contained three mutations (T13P, S15V, V16I) within the N-terminal region corresponding to those found in the slowly desensitizing rP2X2 receptor (Hausmann, Bahrenberg et al. 2014, Mansoor, Lu et al. 2016), enabling an ATP-bound open state to be resolved, as opposed to a desensitized state that was otherwise captured without the P2X3_slow_ mutations (Mansoor, Lu et al. 2016). P2X7 receptors are slowly desensitizing and a cytoplasmic ballast below the cytoplasmic cap likely helps to stabilize the cap (McCarthy, Yoshioka et al. 2019).

**Figure 2.**
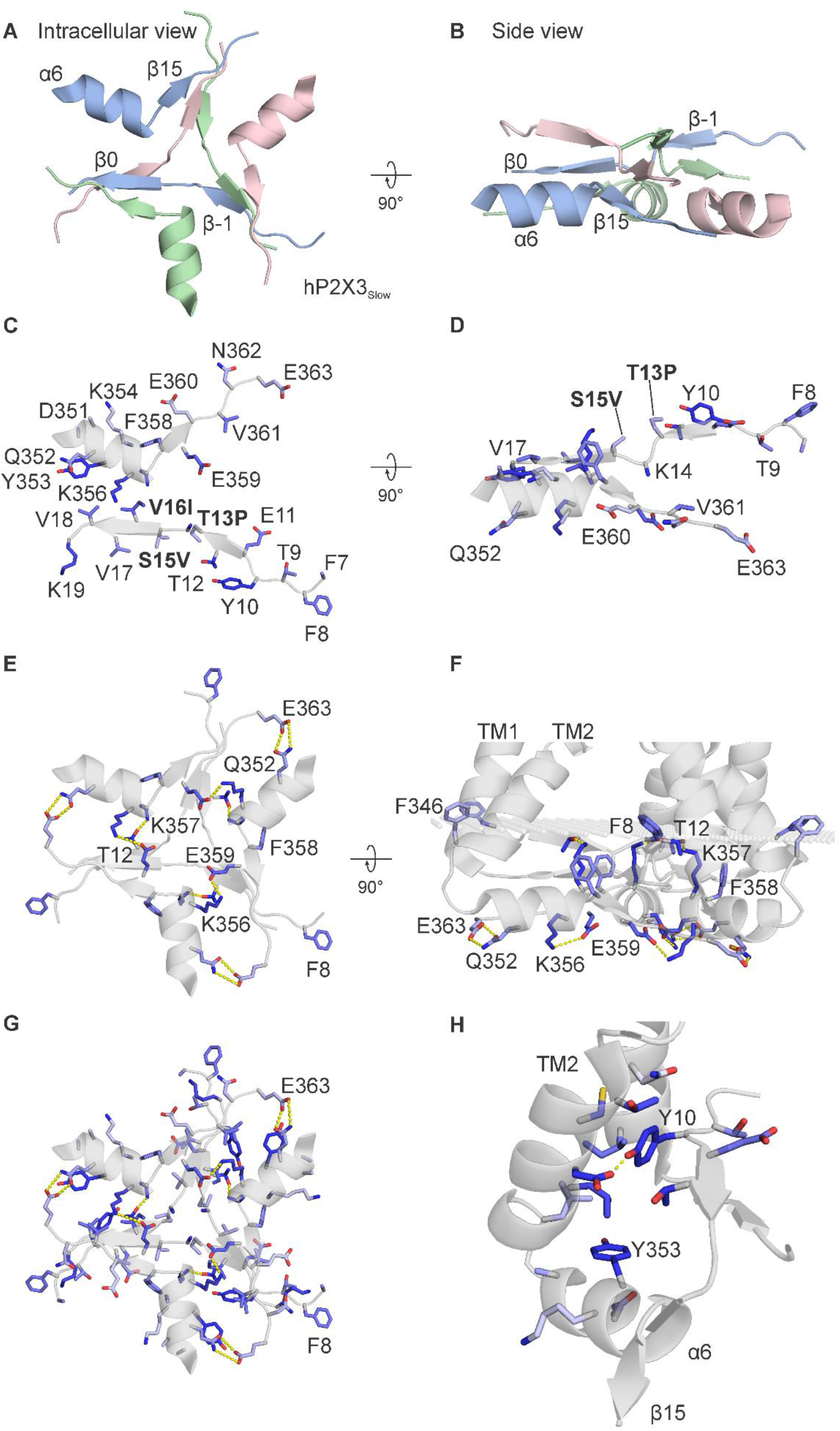
Conservation of residues in the cytoplasmic cap in P2X receptor channels. **(A)** Intracellular view and **(B)** side view of the cytoplasmic cap from hP2X3Slow in complex with ATP (PDB ID: 6ah4) using ribbon representations. **(C)** Intracellular view and **(D)** side view showing a single subunit of hP2X3Slow with side chains shown as sticks and colored according to the alignment quality score calculated from the multiple sequence alignment in Fig. 2 – Fig. Supp. 1, where highly conserved residues are colored in blue and poorly conserved residues are colored in white. Alignment quality score calculated in Jalview based on BLOSUM 62 scores. Residue numbering corresponds to the hP2X3 sequence, and three mutated residues (T13P, S15V, and V16I) that slow desensitization in the construct used for structure determination are highlighted in bold. **(E)** Intracellular view and **(F)** of cytoplasmic cap from hP2X3_Slow_ showing intersubunit side chain interactions between K356 and E359 (3.2-3.8 Å apart); T12 and K357 (2.2-2.3 Å); and Q352 and E363 (2.8-3.8 Å) in yellow. Also pictured are aromatic side chains F8, F346 and F358 facing into the membrane. **(G)** Intracellular view of all three subunits of the cytoplasmic cap of hP2X3_Slow_ with side chains shown as sticks and colored according to the alignment quality score calculated from the multiple sequence alignment in Fig 2 – Fig. Supp. 1. **(H)** View from the lateral fenestration of hP2X3_Slow_ of two conserved tyrosine residues (Y10 and Y353) and the side chains of residues within 4.0 Å. Y10 and D340 of the neighboring subunit are 2.1-2.2 Å apart. The same orientations were used for panels A, C, E, and G and for panels B, D, and F, respectively.

In ATP-bound open state structures of both P2X3_slow_ and P2X7 receptor channels, the cytoplasmic cap would be expected to prevent ions from permeating at the three-fold axis of the channel (Fig. 1D,E; Fig. 2 – Fig Supp 1). While the intracellular lateral fenestrations formed by the TM helices and the cytoplasmic cap are substantial in size, the presence of a lipid membrane would likely diminish the water accessible surface of the fenestrations considerably (Fig. 1A,B). We began by using the OPM server (Lomize, Pogozheva et al. 2012) to predict the boundaries of the lipid membrane, which suggests that the intracellular lateral fenestrations are largely buried within the membrane (Fig. 1B; membrane boundaries depicted with planes of gray dots). Many of the residues lining the edges of the lateral fenestrations are hydrophobic in P2X receptors (Fig. 1B, right), consistent with this structural element residing mostly within the lipid membrane.

To explore whether the cytoplasmic cap is a conserved structural element in P2X receptor channels, we constructed a multiple sequence alignment using all seven subtypes and examined the extent to which residues forming the cytoplasmic cap are conserved (Fig. 2; Fig. 2 – Figure Supp. 1A; Fig. 2 – Figure Supp. 2). Overall, residues within the three ß-strands (ß-1, ß0 and ß15) and one α-helix (α6) forming the cytoplasmic cap are remarkably well-conserved (Fig. 2C; Fig. 2 – Figure Supp. 1A). In the structure of hP2X3Slow with ATP bound, three pairs of residues form hydrogen bonds, including K356 and E359, T12 and K357 and Q352 and E363 (Fig. 2E,F). In addition, although the side chain of K14 was not well-resolved, it is positioned nearby to E359 and is another candidate for participating in stabilizing hydrogen bonds within the cap. All of these polar residue interactions occur at subunit interfaces, and most of the participating residues are well-conserved across all subtypes of P2X receptors (Fig. 2E,F; Fig. 2 – Fig. Supp. 1A,C). The cytoplasmic cap also contains two conserved Tyr residues (Y10 and Y353), each of which are surrounded by hydrophobic residues likely to contribute to stabilizing hydrophobic interactions (Fig. 2H). In the case of Y10, these interactions occur at subunit interfaces, whereas in the case of Y353 the interactions are within each individual subunit, and again these interactions are well-conserved across different P2X receptors (Fig. 2H; Fig. 2 – Fig. Supp. 1A,D). Y10 also forms a well-conserved intersubunit hydrogen bond interaction with D340, located at the base of the TM2 helix of the neighboring subunit (Fig. 2H; Fig.2 – Fig. Supp. 1A,D). We also identified a series of additional aromatic residues (F8, F346 and F358) positioned near to where the OPM server would position the polar headgroup region of the inner leaflet of the membrane, and again these are well-conserved across different subtypes of P2X receptors (Fig. 2F). All of these stabilizing interactions observed in the cytoplasmic cap of hP2X3Slow are also seen in the structure of hP2X7 (Fig. 2 – Fig. Supp. 1A-D) and the participating residues are conserved in most subtypes of P2X receptors (Fig. 2 – Fig. Supp. 1A), suggesting that structural elements related to the caps seen in P2X3 and P2X7 are likely present in all P2X receptor channels.

In studying the structure of the cytoplasmic cap and intracellular lateral fenestrations in hP2X3_Slow_, we noticed the presence of a Glu residue (E11; Fig. 1A,B, D) at the intracellular edge of the lateral fenestration that is conserved in most P2X receptor channels (Fig. 1C), with the exception of both P2X5 and P2X6, which contain a Lys at this position. Although P2X6 channels do not express as homomeric channels (Le, Babinski et al. 1998) and therefore their permeation properties are poorly understood, homomeric P2X5 receptors can be expressed and have been reported to have a significant Cl^-^ permeability (Ruppelt, Ma et al. 2001, Bo, Jiang et al. 2003). We therefore thought this position in P2X receptors might provide a foothold to begin interrogating whether ions permeate through the lateral fenestrations and whether residues in this region influence ion selectivity.

### Accessibility of a conserved Glu in the lateral fenestrations of P2X2 receptor channels

To explore whether ions might permeate through the lateral fenestrations, we decided to investigate the accessibility of an introduced Cys residue to thiol reactive methanethiosulfonate (MTS) compounds. The rP2X2 receptor channel has been extensively used for accessibility studies because it slowly desensitizes and therefore is well-suited for examining changes in accessibility between open and closed states (Li, Chang et al. 2008, Li, Kawate et al. 2010, Kawate, Robertson et al. 2011, Heymann, Dai et al. 2013). In addition, the rP2X2-3T construct, wherein three native Cys residues were substituted with Thr residues, is insensitive to MTS compounds unless additional Cys residues are introduced and has been extensively used for accessibility studies (Li, Chang et al. 2008, Li, Kawate et al. 2010, Kawate, Robertson et al. 2011, Heymann, Dai et al. 2013). We began by introducing a Cys into rP2X2-3T at the position equivalent E11 in hP2X3 (E17 in the rP2X2), reasoning that if this acidic residue lines the lateral fenestrations in P2X2 receptor channels, introducing a Cys at this position would be accessible to thiol-reactive compounds. The concentration-dependence for activation of the E17C mutation in rP2X2-3T by free ATP is similar to rP2X2-3T (Fig. 3 – Fig. Supp. 1), suggesting that the mutant does not dramatically alter the gating mechanism of the receptor. We initially explored the accessibility of 2-trimethylaminoethyl methanethiosulfonate (MTSET) applied from the extracellular side of the membrane to E17C, either when channels are open in the presence of ATP (Fig. 3A,C), or when they are closed in the absence of ATP (Fig. 3B,D). Since MTSET has a fixed positive charge at neutral pH, it cannot cross the membrane and only reacts when a Cys is positioned within an aqueous environment within the channel protein (Holmgren, Liu et al. 1996). Consistent with previous studies (Li, Chang et al. 2008), when we applied MTSET to the external solution for cells expressing the background rP2X2-3T construct, we observed no discernible effect of the reagent when applied in the absence or presence of ATP (Fig. 3A,B,E,F). In contrast, application of MTSET in the presence of ATP resulted in robust and irreversible inhibition of the E17C rP2X2-3T channel (Fig. 3C,E). Application of external MTSET in the absence of ATP had no discernible effect on subsequent current activation by external ATP (Fig. 3D,F), suggesting that the MTS reagent can only access E17C through the ion permeation pathway from the extracellular side of the membrane when the channel is open. To confirm that MTSET accessibility to E17C requires activation by ATP, we confirmed that application of MTSET in the presence of ATP produced robust inhibition following application of the MTSET in the absence of ATP (Fig. 3D,F). From these results, we conclude that a Cys residue introduced at E17 within the intracellular lateral fenestrations is accessible to MTSET applied from the external solution when the pore of the channel has been opened with external ATP.

**Figure 3.**
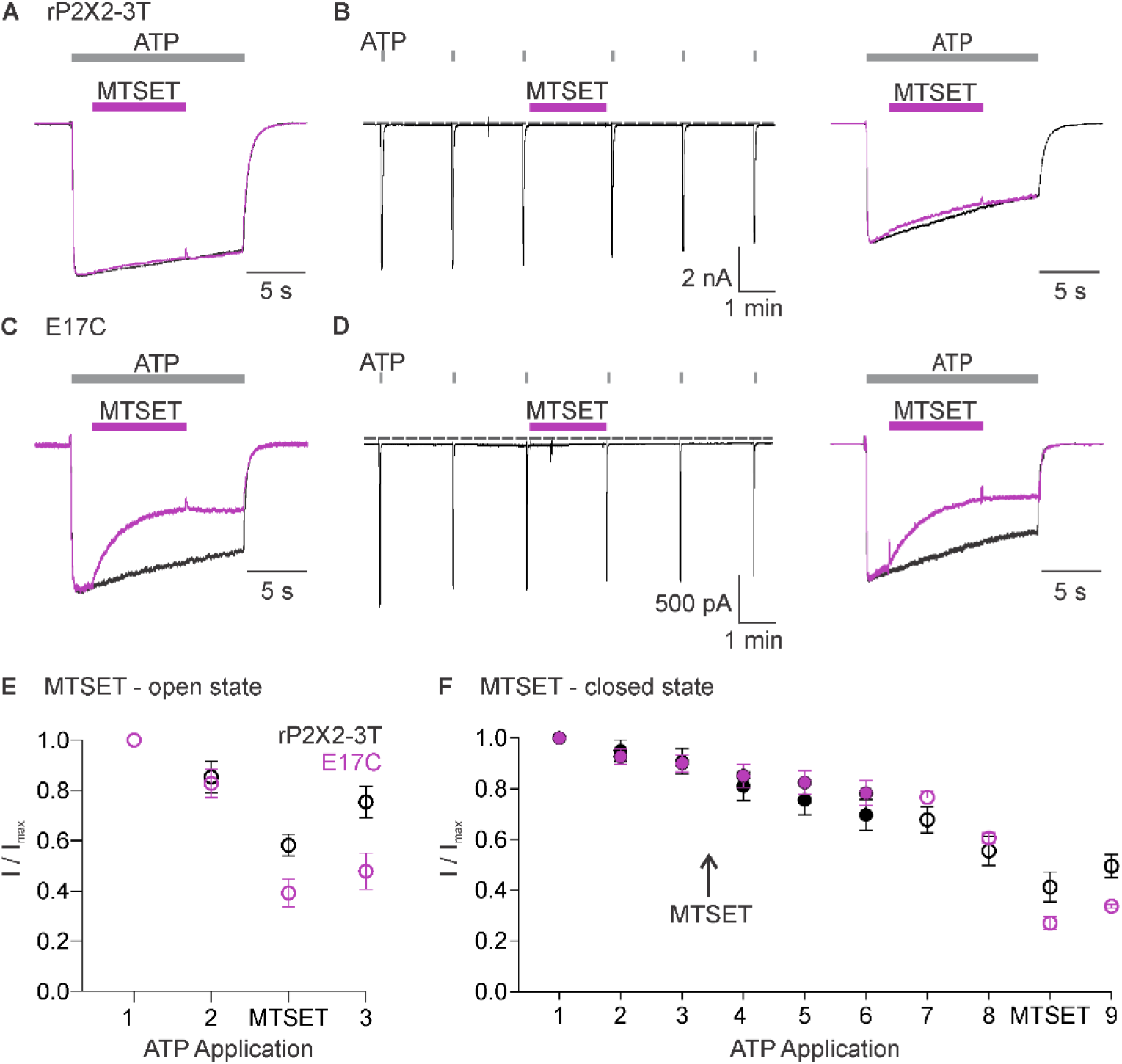
Accessibility E17C in rP2X2-3T to extracellular MTSET (5 inch) **(A)** Testing for modification of rP2X2-3T by extracellular MTSET after opening channels with extracellular ATP. Consecutive current traces elicited by extracellular ATP application without (black trace) or with subsequent application of 1 mM extracellular MTSET (purple trace). ATP (1 μM) was applied for 15 sec at 2 min intervals three times with MTSET application only during the second ATP application. Superimposed and scaled current traces are shown for the first and second application of ATP. **(B)** Testing for modification of rP2X2-3T by extracellular MTSET when channels are closed. ATP (1 μM) was applied six times for 2 s at 2 min intervals and MTSET (1 mM) was applied in the absence of ATP after the third ATP application. Following application of MTSET in the closed state, MTSET was applied again in the presence of ATP to the same cell to serve as a control, using the same protocol described in A. **(C)** Testing for modification of E17C rP2X2-3T by extracellular MTSET after opening channels with extracellular ATP. Consecutive current traces elicited by extracellular ATP application without (black trace) or with subsequent application of 1 mM extracellular MTSET (purple trace). Same protocol as in A. **(D)** Testing for modification of E17C rP2X2-3T by extracellular MTSET when channels are closed. Same protocol as in B. **(E)** Normalized ATP current amplitudes for MTSET application in the presence of ATP for rP2X2-3T (black symbols; n=3) and E17C rP2X2-3T (purple symbols; n=5) using the protocols illustrated in panels A and C. ATP application 1 corresponds to the peak current amplitude for the first control ATP application within 1 s of applying ATP, 2 corresponds to the peak current amplitude immediately before MTS application for the second ATP application, measured within 1 s of ATP application. The ATP application labeled as MTSET corresponds to the current amplitude 10 s into the second ATP application at the end of the MTS application. 3 corresponds to the peak current amplitude during the third ATP application measured within 1 s of applying ATP. **(F)** Normalized ATP current amplitudes during MTSET application in the absence of ATP. Filled black circles (n=3) are from experiments with rP2X2-3T and filled purple circles (n=4) are from experiments with E17C rP2X2-3T) using the protocols illustrated in panels B and D. MTSET was applied in the absence of ATP between ATP application 3 and 4 as indicated. Open symbols are ATP current amplitudes when testing for open state modification by MTSET after testing for closed state modification. Open black circles (n=3) for rP2X2-3T and open purple circles (n=3) for E17C rP2X2-3T using the protocols illustrated on the right in panels B and D. For open state modification following closed state modification, application 7 is a control ATP application, and MTSET was applied during application 8 along with ATP. Data point 8 is the initial peak current immediately before MTSET was applied and the subsequent data point is at the end of the MTSET application in the presence of ATP after the reagent has had time to react, as illustrated in the right panel of B and D. Data shown in E and F are mean ± SEM, some error bars are smaller than symbols.

We next studied modification by 2-aminoethyl methanethiosulfonate (MTSEA), which unlike MTSET, can readily cross membranes (Holmgren, Liu et al. 1996) and therefore provides the means to assess the accessibility of E17C from the intracellular side of the membrane. Similar to MTSET, external application of MTSEA was without discernible effect on rP2X2-3T (Fig. 4, A,B,E,F). Also similar to MTSET, external application of MTSEA in the presence of ATP produced robust and irreversible inhibition of ATP-activated current for the E17C rP2X2-3T channel (Fig. 4C,D,E), suggesting that external MTSEA can access E17 through the ion permeation pathway. However, in contrast to MTSET, external application of MTSEA in the absence of ATP produced discernible inhibition of currents activated by subsequent application of ATP (Fig. 4D,F). Inhibition produced by external application of MTSEA in the absence of ATP prevented subsequent inhibition by external application of MTSEA in the presence of ATP (Fig. 4D,F), suggesting that external MTSEA can access E17C through the ion permeation pathway when the channel is open, or cross the membrane and react with E17C from the intracellular side when the channel is closed.

**Figure 4.**
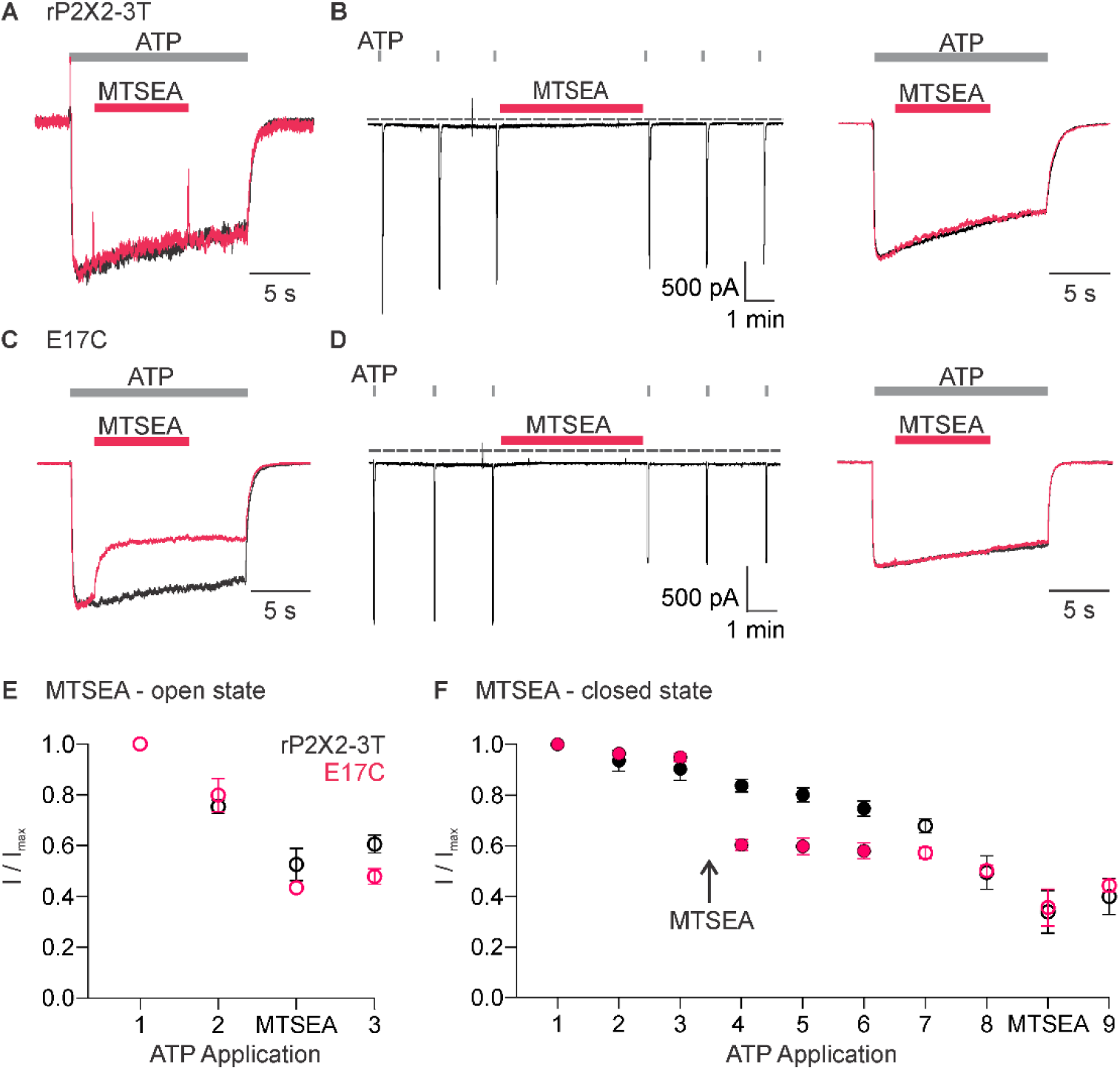
Accessibility E17C in rP2X2-3T to extracellular MTSEA (5 inch) **(A)** Testing for modification of rP2X2-3T by extracellular MTSEA after opening channels with extracellular ATP. Consecutive current traces elicited by extracellular ATP application without (black trace) or with subsequent application of 1 mM extracellular MTSEA (red trace). ATP (1 μM) was applied for 15 sec at 2 min intervals three times with MTSEA application only during the second ATP application. Superimposed and scaled current traces are shown for the first and second application. **(B)** Testing for modification of rP2X2-3T by extracellular MTSEA when channels are closed. ATP (1 μM) was applied six times for 2 s at 2 min intervals and MTSEA (1 mM) was applied in the absence of ATP after the third ATP application. Following application of MTSEA in the closed state, MTSEA was applied again in the presence of ATP to the same cell to serve as a control. **(C)** Testing for modification of E17C rP2X2-3T by extracellular MTSEA after opening channels with extracellular ATP. Current traces elicited by extracellular ATP application without (black trace) or with subsequent application of 0.5 mM extracellular MTSEA (red trace). Same protocol as in A. **(D)** Testing for modification of E17C rP2X2-3T by extracellular MTSEA when channels are closed. Same protocol as in B. **(E)** Normalized ATP current amplitudes for MTSEA application in the presence of ATP for rP2X2-3T (black symbols; n=5) and E17C rP2X2-3T (red symbols; n=4) using the protocols illustrated in panels A and C. ATP application 1 corresponds to the peak current amplitude for the first control ATP application, 2 corresponds to the peak current amplitude immediately before MTS application for the second ATP application, measured within 1 s of ATP application. The ATP application labeled MTSEA corresponds to the current amplitude 10 s into the ATP application and at the end of the MTSEA application during the second ATP application. 3 corresponds to peak current amplitude 2 min after MTS application during the third ATP application, measured within 1 s of ATP application. **(F)** Normalized ATP current amplitudes during MTSEA application in the absence of ATP. Filled black circles (n=6) are from experiments with rP2X2-3T and filled red circles (n=4) are from experiments with E17C rP2X2-3T) using the protocols illustrated in panels B and D. MTSEA was applied in the absence of ATP between ATP application 3 and 4 as indicated. Open symbols are ATP current amplitudes when testing for open state modification by MTSEA after testing for closed state modification. Open black circles (n=3) for rP2X2-3T and open red circles (n=3) for E17C rP2X2-3T using the protocols illustrated in panels B and D. For open state modification following closed state modification, application 7 is a control ATP application, and MTSEA was applied during application 8 along with ATP. Data point 8 is the initial peak current before MTSEA was applied and the subsequent data point is at the end of the MTSEA application in the presence of ATP after the reagent has had time to react, as illustrated in the right panel of B and D. Data shown in E and F are mean ± SEM, some error bars are smaller than symbols.

Having observed robust current inhibition by both MTSET and MTSEA, we next measured the rates of modification of E17C by MTSET and MTSEA in the presence of ATP and compared them with those measured in previous studies where rates of modification were reported for Cys substitutions within the pore-lining TM2 helix (Li, Chang et al. 2008). The time courses for MTS modification in the presence of ATP could be reasonably well fit by a single exponential function (Fig. 5A,B), from which we calculate that MTSET modified E17C with a rate of 373 M^-1^s^-1^ (n = 5), while the smaller MTSEA modified that position with a rate of 2565 M-^1^s^-1^ (n = 4). From previous MTSET modification rates obtained in the presence of ATP for multiple positions within the TM2 helix, we can appreciate that rates of modification diminish as Cys residues are introduced deeper into the pore (Fig. 5D). Notably, the modification rate for externally applied MTSET at E17C is only incrementally slower than the deepest position in the transmembrane region when applying the reagent to the external solution (Fig. 5D). Given that the reagents must traverse the entire TM pore to react with E17C, these relatively rapid rates of modification suggest that E17C is positioned in a high dielectric environment that is favorable for modification. Although lipid molecules likely occupy the upper regions of the lateral fenestration, it seems unavoidable that the most intracellular end of the lateral fenestrations where E17 is located is an aqueous environment. From these experiments examining the modification of E17C within the lateral fenestrations, we conclude that this residue lines the intracellular end of the ion permeation pathway.

**Figure 5.**
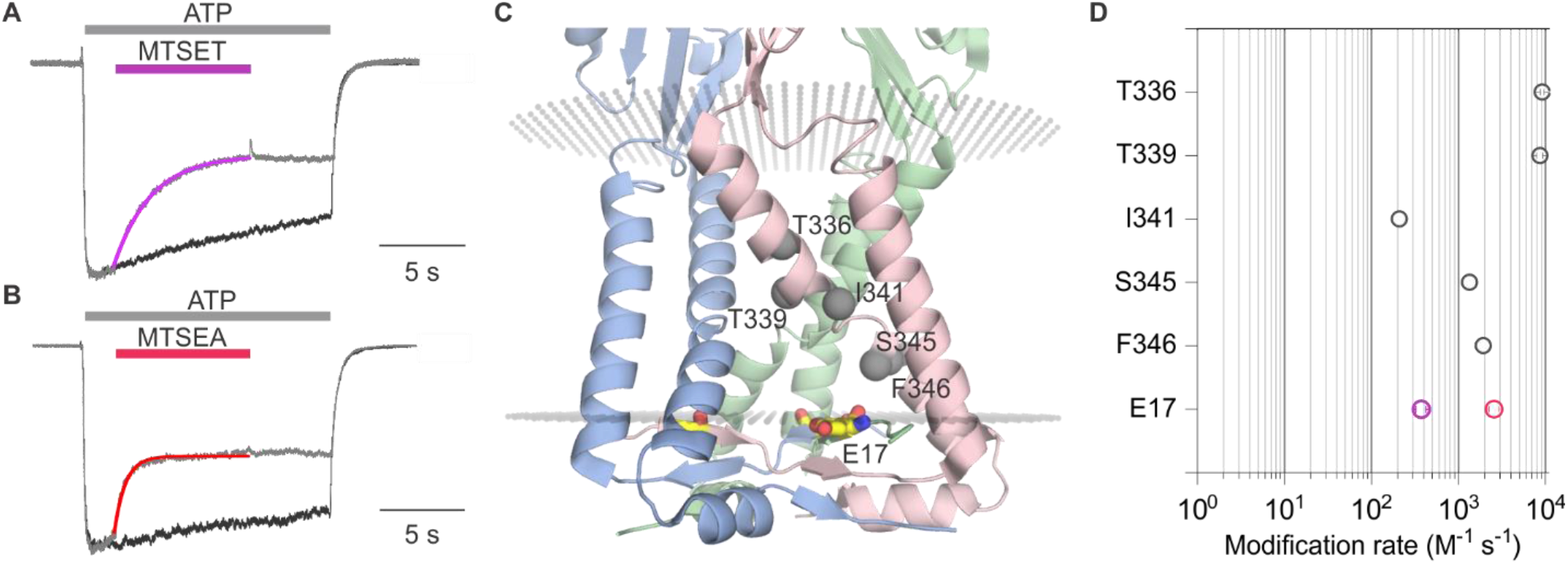
Rates of MTS Modification in rP2X2 for E17C and comparison with rates for the pore-lining TM2 helix (7 inch) **(A)** Fit of a single exponential function (purple relation) to the time course of current inhibition by MTSET (1 mM) for E17C rP2X2-3T in the presence of ATP (1 μM). **(B)** Fit of a single exponential function (red relation) to the time course of current inhibition by MTSEA (0.5 mM) for E17C rP2X2-3T in the presence of ATP (1 μM). **(C)** Side view of the transmembrane helices and cytoplasmic cap from the structure of hP2X3_Slow_ with ATP bound (PDB ID: 6ah4). MTS accessible residues along TM2 are labeled and each residue alpha carbon is represented as a gray sphere. **(D)** Comparison of MTS modification rates measured in the presence of ATP for E17C rP2X2-3T (purple open circles for MTSET and red open circles for MTSEA) with MTSET reactive residues in TM2 (gray open circles) from (Li, Chang et al. 2008). Data is mean ± SEM, some error bars are smaller than symbols.

### Influence of lateral fenestrations on cation selectivity of P2X2 receptor channels

Having uncovered evidence that E17 lines the ion permeation pathway, we next explored whether this conserved acidic residue is a determinant of the cation selectivity in P2X2 receptors. Although P2X receptor channels are permeable to Na^+^ and to varying extents Ca^2+^ (Migita, Haines et al. 2001, Samways and Egan 2007, Samways, Li et al. 2014), we are unaware of attempts to measure the relative permeability of cations to anions for most subtypes, except for those that have noted a measurable anion permeability for P2X5 receptor channels (Ruppelt, Ma et al. 2001, Bo, Jiang et al. 2003). We began by assessing the relative permeability of wild-type rP2X2 to cations over anions by initially measuring equilibrium reversal potentials (V_rev_) using ramps and switching between symmetrical solutions containing 140 mM NaCl on both the external and internal sides and asymmetrical solutions containing 140 mM NaCl internally and 40 mM NaCl externally, using either sucrose or glucose to maintain osmolarity (Fig. 6A). To minimize errors due to possible ion accumulation on the intracellular side (Li, Toombes et al. 2015), we maintained the cell in symmetrical solutions and only assessed the impact of lowering external NaCl with ATP applications just long enough to measure current-voltage (I-V) relations and determine V_rev_. For wild-type rP2X2 receptor channels, we measured V_rev_ with symmetric solutions near 0 mV and upon switching to the external containing 40 mM NaCl observed that V_rev_ shifted to around −32 mV (Fig. 6A,G), from which we calculate a pNa^+^:pCl^-^ of 20:1 using the Goldman-Hodgkin-Katz equation (see Materials and Methods), indicative of a strong preference for cations over anions. Considering that rP2X2 is inwardly rectifying and thus passes outward currents less favorably than inward currents (Zhou and Hume 1998, Fujiwara and Kubo 2004, George, Swartz et al. 2019), we also used voltage step protocols and measured the initial tail current on stepping from −60 mV to more positive voltages, enabling the measurement of relatively linear instantaneous I-V relations and an independent determination of V_rev_ (Fig. 6B,C). As with the ramp protocol, the step protocol resulted in shifts in V_rev_ from near 0 mV to near −32 mV upon switching the external solution from 140 mM NaCl to ones containing 40 mM NaCl (Fig. 6B,C,G). Collectively, these results with either ramp or step protocols establish that rP2X2 exhibits a strong preference for cations over anions.

**Figure 6.**
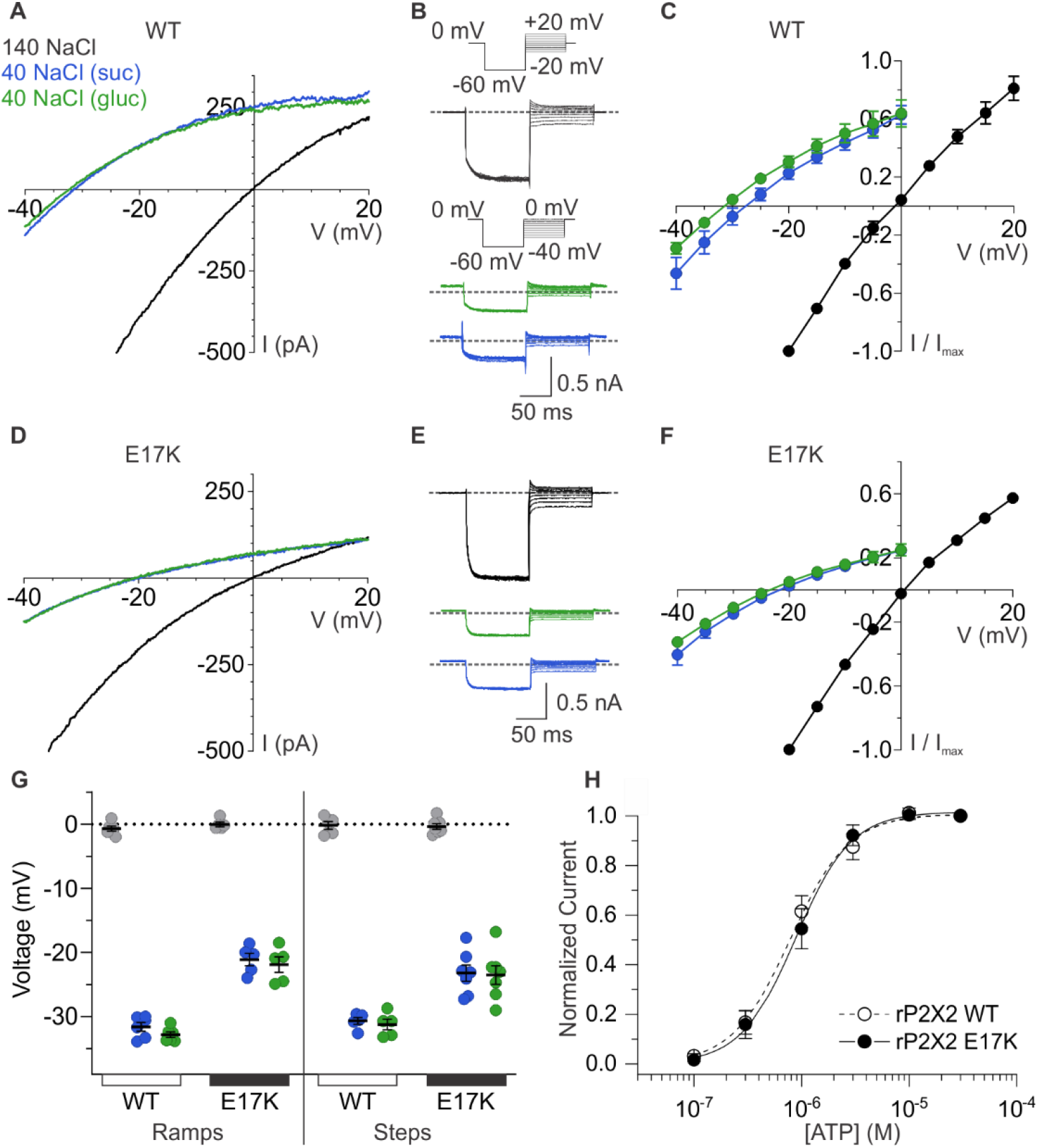
Ion selectivity of rP2X2 and influence of the E17K mutation. (5 inch) **(A)** Representative I-V relationships for WT rP2X2 obtained from the same cell using 0.5 s voltage ramps from −60 mV to +60 mV from a holding potential of −60 mV in symmetric 140 mM NaCl solution (black) and two asymmetric NaCl solutions (internal: 140 mM NaCl; external:40 mM NaCl with glucose in green or 40 mM NaCl with sucrose in blue). Voltage ramps were applied in the absence and presence of 30 μM ATP in each of the three external solutions and currents from ramps recorded in the absence of ATP were subtracted from currents in the presence of ATP to obtain the ATP activated currents shown. **(B)** Representative current traces from a cell expressing WT rP2X2 elicited using voltage step protocols in symmetrical 140 mM NaCl solution and two asymmetrical solutions as in A. The 300 ms voltage step protocols were conducted with a holding potential of 0 mV and step to −60 mV for 100 ms before stepping to a series of voltages in increments of 5 mV. The range of voltages tested for symmetrical solutions was −20 mV to +20 mV and for asymmetrical solutions was −40 mV to 0 mV steps. ATP-activated currents were obtained by subtracting control currents recorded in the absence of ATP from currents collected in the presence of 30 μM ATP. The gray dotted line represents 0 pA. **(C)** Normalized instantaneous I-V relationships for WT rP2X2 obtained using voltage step protocols illustrated in panel B in symmetrical 140 mM NaCl and two asymmetrical solutions. Instantaneous current was measured 0.5-1 ms following the step from −60 mV to the range of voltages indicated. Data points represent the mean steady-state current ± SEM (n=6 for each condition). **(D)** Representative I-V relationships for the E17K mutant of rP2X2 obtained from the same cell in symmetrical 140 mM NaCl and two asymmetric solutions using the voltage ramp protocol from panel A. **(E)** Representative current traces from a cell expressing the E17K mutant of rP2X2 elicited using the same voltage step protocols from panel B in symmetrical 140 mM NaCl and two external solutions containing 40 mM NaCl. **(F)** Normalized instantaneous I-V relationships for the E17K mutant of rP2X2 obtained using voltage step protocols illustrated in panel E in symmetrical 140 mM NaCl and two external solutions containing 40 mM NaCl. Protocols were identical to those in C and data points represent the mean steady-state current ± SEM (n=7). **(G)** Reversal potential measurements for WT and E17K rP2X2 in symmetrical and asymmetric NaCl solutions using either ramp or step protocols. Mean and SEM are indicated with bars and individual measurements are indicated with filled colored circles using the same color code as in A. WT rP2X2 ramp protocol reversal potentials were −0.7 ± 0.4 mV in symmetric solution, −31.6 ± 0.7 mV in asymmetric solution with sucrose, and −32.8 ± 0.4 mV in asymmetric solution with glucose. E17K rP2X2 ramp protocol reversal potentials were 0 ± 0.4 mV in symmetric solution, −21.1 ± 1.0 mV in asymmetric solution with sucrose, and −21.9 ± 1.2 mV in asymmetric solution with glucose. WT rP2X2 step protocol reversal potentials were −0.2 ± 0.6 mV in symmetric solution, −30.7 ± 0.5 mV in asymmetric solution with sucrose, and −31.2 ± 0.8 mV in asymmetric solution with glucose. E17K rP2X2 step protocol reversal potentials were −0.4 ± 0.4 mV in symmetric solution, −23.3 ± 1.3 mV in asymmetric solution with sucrose, and −23.5 ± 1.5 mV in asymmetric solution with glucose. For WT, ramp protocols n=6 and step protocols n=5. For E17K, ramp protocols n=5 and step protocols n=7. **(H)** Normalized concentration-dependence for ATP activation of WT rP2X2 (open circles and dashed line, n=4) and the E17K mutant of rP2X2 (black circles, n=5). Smooth curves are fits of the Hill equation to the data with EC50 and nH values of 0.8 ± 0.1 μM and 1.7 ± 0.1 for WT and 0.9 ± 0.1 μM and 1.6 ± 0.1 for E17K.

To explore whether E17 contributes to the cation selectivity, we mutated this residue to Lys, the residue that occupies the equivalent position in anion permeable P2X5 receptor channels (Fig. 1C). The E17K mutation did not appreciably alter the concentration-dependence for activation of the channel for ATP (Fig. 6H), suggesting that the mutation does not perturb gating. However, the E17K mutation did alter the shift in V_rev_ produced by lowering external NaCl concentration, in this case shifting to between −22 and −25 mV depending on whether we used ramp or step protocols, and whether ions were substituted with sucrose or glucose (Fig. 6D-G). If we calculate the relative permeability of cations to anions, the E17K diminished pNa^+^:pCl^-^ to 10:1 from 20:1 measured for the wild-type receptor. Although these results establish that E17 is not the sole determinant of cation selectivity in P2X2 receptor channels, mutation of this residue detectably alters cation selectivity, consistent with this residue residing within the ion permeation pathway on the intracellular side of the membrane.

### Influence of lateral fenestrations on anion permeability of P2X5 receptor channels

The P2X5 receptor channel has not been extensively studied because heterologous expression results in only small macroscopic currents in response to ATP application (Bo, Jiang et al. 2003, Sun, Liu et al. 2019, Illes, Muller et al. 2021). We initially expressed P2X5 receptor channels from rat and mouse and observed that expression of the mouse clone produced the largest ATP-activated currents in our standard extracellular solution lacking divalent cations (see Materials and Methods). We also observed a robust yet variable sensitization phenomenon whereby ATP-activated currents in cells expressing mP2X5 increased in amplitude over the time course of several minutes (Fig. 7 A,B). Although we have not yet explored the underlying mechanism, we adopted a standard sensitization protocol (Fig. 7A) wherein we applied a saturating concentration of ATP (3 μM) until ATP-activated current reached steady-state before undertaking further experiments. Examination of the concentration-dependence for activation of mP2X5 revealed a relatively high affinity for ATP in the absence of divalent ions, roughly two orders of magnitude higher than wild-type rP2X2 (Fig. 7C).

**Figure 7.**
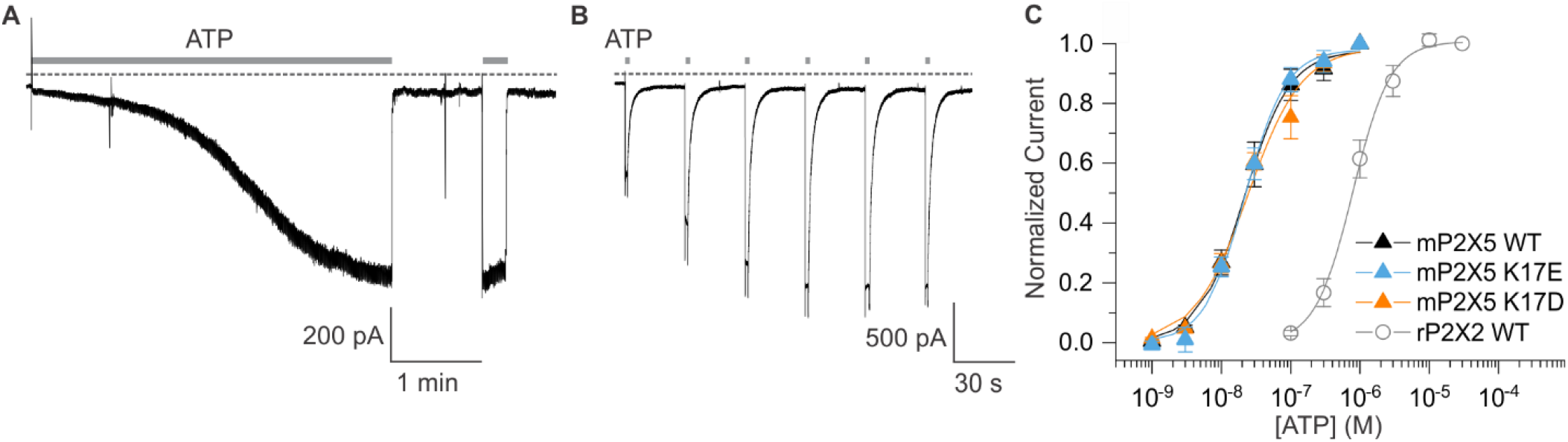
Expression and characterization of mP2X5 (7 inch) **(A)** mP2X5 sensitization upon initial exposure to a sustained external application of ATP (3 μM). Subsequent short ATP applications activate slowly desensitizing currents of similar amplitude compared to that achieved during sensitization. Holding voltage was −60 mV. Gray dotted line represents zero current. **(B)** Sensitization observed while delivering short pulses of saturating ATP (100 μM) given in 30 s intervals. Holding voltage was −60 mV. Gray dotted line represents zero current. **(C)** Concentration-dependence for ATP activation of WT mP2X5 (black filled triangles, n=3), the K17E mutant of mP2X5 (light blue triangles, n=4), and the K17D mutant of mP2X5 (orange triangles, n=3). Smooth curves are fits of the Hill equation to the data with EC_50_ and n_H_ values of 22.5 ± 1.6 nM and 1.3 ± 0.1 for WT, 21.9 ± 1.7 nM and 1.5 ± 0.1 for K17E, and 24.1 ± 4.3 nM and 1.1 ± 0.2 for K17D. Data for WT rP2X2 are shown for comparison from Fig. 6.

We next examined the relative permeability of mP2X5 for cations and anions using the same approach we employed for rP2X2, in this case only using ramps because we observed relatively linear I-V relations for mP2X5 with little if any evidence of inward rectification (Fig. 8A). Remarkably, decreasing the concentration of external NaCl did not shift V_rev_ towards negative voltages, as we observed for rP2X2, but instead produced a slight positive shift in V_rev_ by a few mV (Fig. 8A,D). From these shifts in V_rev_ we calculate that mP2X5 has a slight preference for anions over cations, with a relative permeability of pNa^+^:pCl^-^ of 0.8. These findings confirm earlier reports that P2X5 receptor channels have a much higher Cl^-^ permeability compared to other subtypes of P2X receptor channels (Ruppelt, Ma et al. 2001, Bo, Jiang et al. 2003).

**Figure 8.**
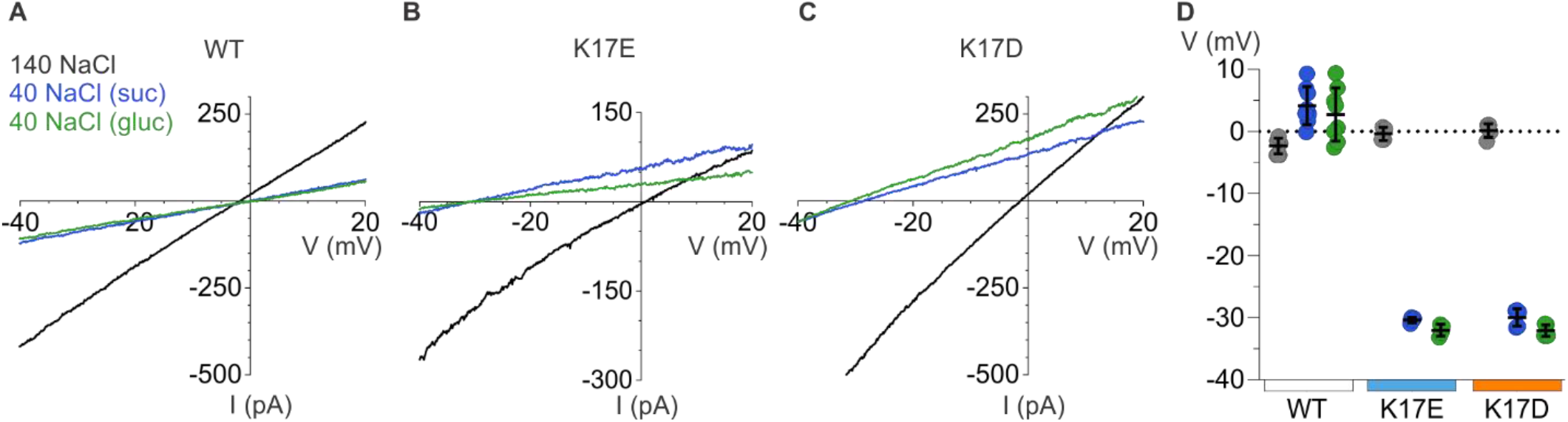
Ion selectivity of mP2X5 and influence of K17E and K17D mutations. (7 inch) **(A)** Representative I-V relationships for WT mP2X5 obtained from the same cell using 0.5 s voltage ramps from −60 mV to +60 mV from a holding potential of −60 mV in symmetric 140 mM NaCl solution (black) and two asymmetric NaCl solutions (internal: 140 mM NaCl; external:40 mM NaCl with glucose in green, or 40 mM NaCl with sucrose in blue). Voltage ramps were applied in the absence and presence of 3 μM ATP in each of the three conditions and currents from ramps recorded in the absence of ATP were subtracted from currents in the presence of ATP to obtain the ATP activated currents shown. **(B)** Representative I-V relationships for the K17E mutant of mP2X5 obtained from the same cell using conditions as in A. **(C)** Representative I-V relationships for the K17D mutant of mP2X5 obtained from the same cell using conditions as in A and B. **(D)** Reversal potential measurements for WT, K17E and K17D mP2X5 in symmetrical and asymmetrical NaCl solutions using ramp protocols. Mean and SEM are indicated with bars and individual measurements are indicated with filled colored circles using the same color code as in A. WT mP2X5 reversal potentials were −2.3 ± 0.4 mV in symmetric solution, 4.1 ± 1.1 mV in asymmetric solution containing sucrose, and 2.8 ± 1.5 mV in asymmetric solution containing glucose. K17E mP2X5 reversal potentials were −0.4 ± 0.5 mV in symmetric solution, −30.4 ± 0.2 mV in asymmetric solution containing sucrose, and −32 ± 0.5 mV in asymmetric solution containing glucose. K17D mP2X5 reversal potentials were 0.2 ± 0.5 mV in symmetric solution, - 30 ± 0.6 mV in asymmetric solution containing sucrose, and −32.1 ± 0.4 mV in asymmetric solution containing glucose. n=8 for WT, n=4 for K17E, and n=5 for K17D.

To explore whether K17 is a determinant of the ion selectivity of mP2X5 receptor channel, we mutated this position to either Glu or Asp. Neither mutation appreciably altered the concentration-dependence for activation of mP2X5 by external ATP (Fig. 7C), suggesting that as for rP2X2, mutations at this position within the lateral fenestrations do not appreciably alter gating. We then proceeded to measure shifts in V_rev_ upon switching external solutions from 140 mM NaCl to 40 mM NaCl, and strikingly observed that for both mutants, lowering the external NaCl concentration produced dramatic shifts of V_rev_ to voltages between −32 and −35 mV (Fig. 8B-D), similar to what we observed for wild-type rP2X2 receptor channels (Fig. 6A-C,G). From these shifts in V_rev_ we calculate that K17E and K17D mutants are strongly cation selective, with pNa^+^:pCl^-^ of around 20:1, indistinguishable from what we measured for wild-type rP2X2 receptor channels. These findings indicate that K17 is a critical determinant of the unusual anion permeability of mP2X5 receptor channels. Collectively, our results support the idea that the lateral fenestrations observed in recent structures of P2X3 and P2X7 receptor channels contribute to the intracellular end of the ion permeation pathway.

## DISCUSSION

The objective of the present study was to explore whether ion permeation through the internal pore of P2X receptor channels could occur through the lateral fenestrations seen in recent structures of P2X3 and P2X7 receptors. One enigmatic feature of the lateral fenestrations is that they were observed for detergent solubilized proteins, and one might expect that they would be partially or completely buried within the lipid bilayer when the protein resides in a membrane environment, in agreement with many residues at the edges of the lateral fenestrations having hydrophobic side chains (Fig. 1). However, an acidic residue Glu at the intracellular edge of the lateral fenestrations is conserved in P2X receptors thought to be cation selective but is substituted with the basic residue Lys in P2X5 (Fig. 1), the one subtype of P2X receptor reported to also be anion permeable. This correlation between side chain charge and ion selectivity suggested that this residue might face the ion conducting pathway and contribute directly to ion selectivity. For the E17C mutant in rP2X2 receptors, we found that the positively charged MTSET could readily react with this position, but only when the channel was opened by external ATP (Fig. 3). In contrast, the membrane permeant MTSEA could react with E17C either when the channel was opened by external ATP, or when applied in the closed state in the absence of ATP (Fig. 4), indicating that E17C is accessible from both sides of the membrane. Although approximations of P2X receptor channels in the membrane by databases such as OPM suggest that much of the lateral fenestrations are buried in the membrane (Fig. 1), the efficient reaction between MTS and E17C (Fig. 5) suggests that this residue resides in an aqueous environment and is not buried in the membrane, supporting its location within the ion permeation pathway at the intracellular end of the pore. Importantly, charge reversing mutations in rP2X2 and mP2X5 clearly alter the relative permeability of the two channels (Figs. 6 and 8). Although the effect of the mutation in rP2X2 is unambiguous (Fig. 6), the result also suggests that other determinants contribute to cation selectivity for this subtype. In stark contrast, the influence of the mutations in mP2X5 (Fig. 8) suggest that K17 is a major determinant of anion permeability in that subtype. Viewed collectively, our results support the idea that ion permeation occurs through the lateral fenestrations and that these structural elements play important roles in determining the ion selectivity of P2X receptor channels.

An important question remains about whether the cytoplasmic caps seen in structures of slowly desensitizing hP2X3_Slow_ receptors in an ATP-bound open state (Mansoor, Lu et al. 2016) and those observed in both closed and ATP-bound open states of hP2X7 receptors (McCarthy, Yoshioka et al. 2019) are present in other P2X receptor subtypes. Many residues at critical positions in the cytoplasmic cap are conserved in other P2X receptor subtypes (Fig. 2; Fig. 2 – Fig. Supp. 1), supporting the notion that the cap is a conserved feature of P2X receptor channels. Given that the structures of hP2X3_slow_ where the cap is seen contain three mutations corresponding to residues present in the slowly desensitizing P2X2 receptor, it would seem likely that the cap will also be present in P2X2 receptors. Our results with MTS accessibility and ion selectivity experiments in P2X2 are consistent with the structure of P2X3_slow_ containing the cytoplasmic cap (Hausmann, Bahrenberg et al. 2014, Mansoor, Lu et al. 2016), further supporting the presence of a cap in P2X2 receptors. Similarly, the critical influence of K17 in P2X5 on ion selectivity in that subtype would be consistent with the presence of cytoplasmic cap and lateral fenestrations like those present in the P2X3_slow_ structure. Although earlier studies had implicated residues in the outer pore as critical determinants of Ca^2+^ permeability in P2X receptors (Migita, Haines et al. 2001, Samways and Egan 2007, Samways, Li et al. 2014), a recent study also implicated residues in the N-terminus of rP2X7 as determinants of Ca^2+^ permeability (Liang, Samways et al. 2019). Indeed, the E17A mutation in rP2X2 and the equivalent E14A mutation in rP2X7 both diminish fractional Ca^2+^ current (Liang, Samways et al. 2019), supporting a role of the lateral fenestrations in ion permeation and selectivity. In the future, it would be important to solve structures of other P2X receptor subtypes to confirm that the cytoplasmic cap and lateral fenestrations are feature common to all subtypes, and to determine whether the cap is a transient structure, as proposed for P2X3 receptors (Mansoor, Lu et al. 2016) or a more stable one as proposed for P2X7 (McCarthy, Yoshioka et al. 2019). While the dimensions of the TM domain of P2X receptors would suggest the intracellular lateral fenestrations are largely buried within the membrane, it will be interesting to solve structures of these channels in a membrane environment to better understand how lipids might organize around the fenestrations to effectively distort the structure of the membrane to enable efficient permeation of ions through the internal end of the pore.

It will also be fascinating to explore how the ion selectivity of P2X receptor channel subtypes has been tuned to underlie distinct physiological functions in different cellular environments. Although many important roles of cation selective subtype of P2X receptors are emerging, the functions of the anion permeable P2X5 receptor remain largely enigmatic (Khakh and North 2006, Surprenant and North 2009, Schmid and Evans 2019). P2X5 is thought to be widely expressed within neurons in the central nervous system (Guo, Xu et al. 2008) as well as in cardiac and skeletal muscle (Garcia-Guzman, Soto et al. 1996, Ryten, Hoebertz et al. 2001), yet what roles this channel plays will require further investigation. The unique ion selectivity of P2X5 receptors, determined by one critical residue within the lateral fenestrations, suggests that it may play very different roles than other P2X receptor subtypes.

## MATERIALS AND METHODS

### Channel Constructs

Rat P2X2 (rP2X2) (Brake, Wagenbach et al. 1994) cDNA in pcDNA1 was generously provided by Dr. David Julius (University of California, San Francisco, CA). Mouse P2X5 (mP2X5) cDNA in pCMV6-Entry was purchased from OriGene and subcloned into pcDNA3.1. A previously characterized rP2X2 construct where three native cysteines (C9, C348 and C430) were mutated to threonine (rP2X2-3T) was used as a background construct for MTS experiments because it is insensitive to MTS reagents yet displays functional properties that are similar to the wild-type channel (Li, Chang et al. 2008). All mutations in rP2X2 and mP2X5 were made using the QuikChange Lightning technique (Agilent Technologies) and confirmed by DNA sequencing (Macrogen).

### Cell Culture

Human Embryonic Kidney (HEK293) cells were cultured in Dulbecco’s modified Eagle’s medium (DMEM) supplemented with 10% fetal bovine serum (FBS) and 10 mg L^-1^ of gentamicin. HEK293 cells between passage numbers 5-20 were used and passaged when cells were between 40-80% confluent. The cells were treated with trypsin and then seeded on glass coverslips at about 15% of the original confluency in 35 mm petri dishes. Transfections were done using the FuGENE6 Transfection Reagent (Promega). Transfected cells were incubated at 37°C with 95% air and 5% CO2 overnight for use in whole-cell recordings; 16-24h for reversal potential measurements and 40-48h for MTS experiments. All P2X constructs were co-transfected with a green fluorescent protein cDNA to identify transfected cells.

### Electrophysiology

Membrane currents were recorded from HEK293 cells using the whole-cell patch-clamp technique. When the whole-cell configuration was established, the desired extracellular solution was continuously perfused onto the cell through a gravity-fed perfusion system. Membrane voltage was controlled using an Axopatch 200B patch-clamp amplifier (Axon Instruments) and currents digitized using a Digidata 1440A interface board and pCLAMP 10 software. Membrane currents were collected with a sampling rate of 10 kHz and filtered at 2 kHz. The standard extracellular solution used to form GΩ seals contained 140 mM NaCl, 5.4 mM KCl, 2 mM CaCl2, 0.5 mM MgCl2, 10 mM HEPES, and 10 mM D-glucose, adjusted to pH 7.3 with 1M NaOH, with an osmolality of about 300 mmol/kg. The standard pipette solution contained 140 mM NaCl, 10 mM HEPES, and 10 mM EGTA, adjusted to pH 7.0 with 1M NaOH. The standard extracellular recording solution used for obtaining concentration-response relations and MTS accessibility experiments contained 140 mM NaCl and 10 mM HEPES, adjusted to pH 7.3 with 1M NaOH, with an osmolality of about 275 mmol/kg. Reversal potential measurements were made using the standard extracellular solution and one containing 40 mM NaCl, 10 HEPES, and 175 mM D-glucose or sucrose, adjusted to pH 7.3 with 1M NaOH, with an osmolality of about 270 mmol/kg. Bath and ground chambers were connected by an agar bridge containing 3M KCl. Liquid junction potentials between internal and external solutions were measured and found to be within ± 1.5 mV and all voltages were not corrected. Solutions containing ATP were freshly prepared for use on the same day. Stock solutions of 100 mM MTS reagents (bromide salt; Toronto Research Chemicals) were prepared daily in deionized water and stored on ice. MTS reagents were diluted to the desired concentration less than 2 minutes prior to application of the reagent to individual cells.

### Data Analysis

Concentration-response relationships for ATP were obtained for each mutant channel and the Hill equation fit to the data according to:

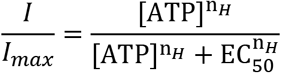

where *I* is the normalized current at a given ATP concentration, *I_max_* is the maximum normalized current, EC50 is the concentration of ATP ([ATP]) that elicits half-maximal currents, and nH is the Hill coefficient.

Time constants for MTS modification (τ) were obtained by fitting the relaxation of current inhibition by MTS reagents with a single exponential function (Fig. 4) and modification rates *(R)* were calculated according to:

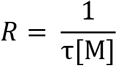

where [M] is the concentration of the MTS reagent.

The relative permeability of P_Cl_/P_Na_ was estimated using the Goldman-Hodgkin-Katz equation (Hille 2001):

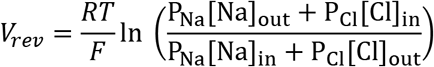

## Acknowledgements

We thank Angela Ballesteros, Surbhi Dhingra and members of the Swartz laboratory for helpful discussions. This research was supported by the Intramural Research Program of the National Institute of Neurological Disorders and Stroke, NIH, Bethesda, MD to KJS (NS003018).

---

**Figure 2 – Figure Supplement 1.**
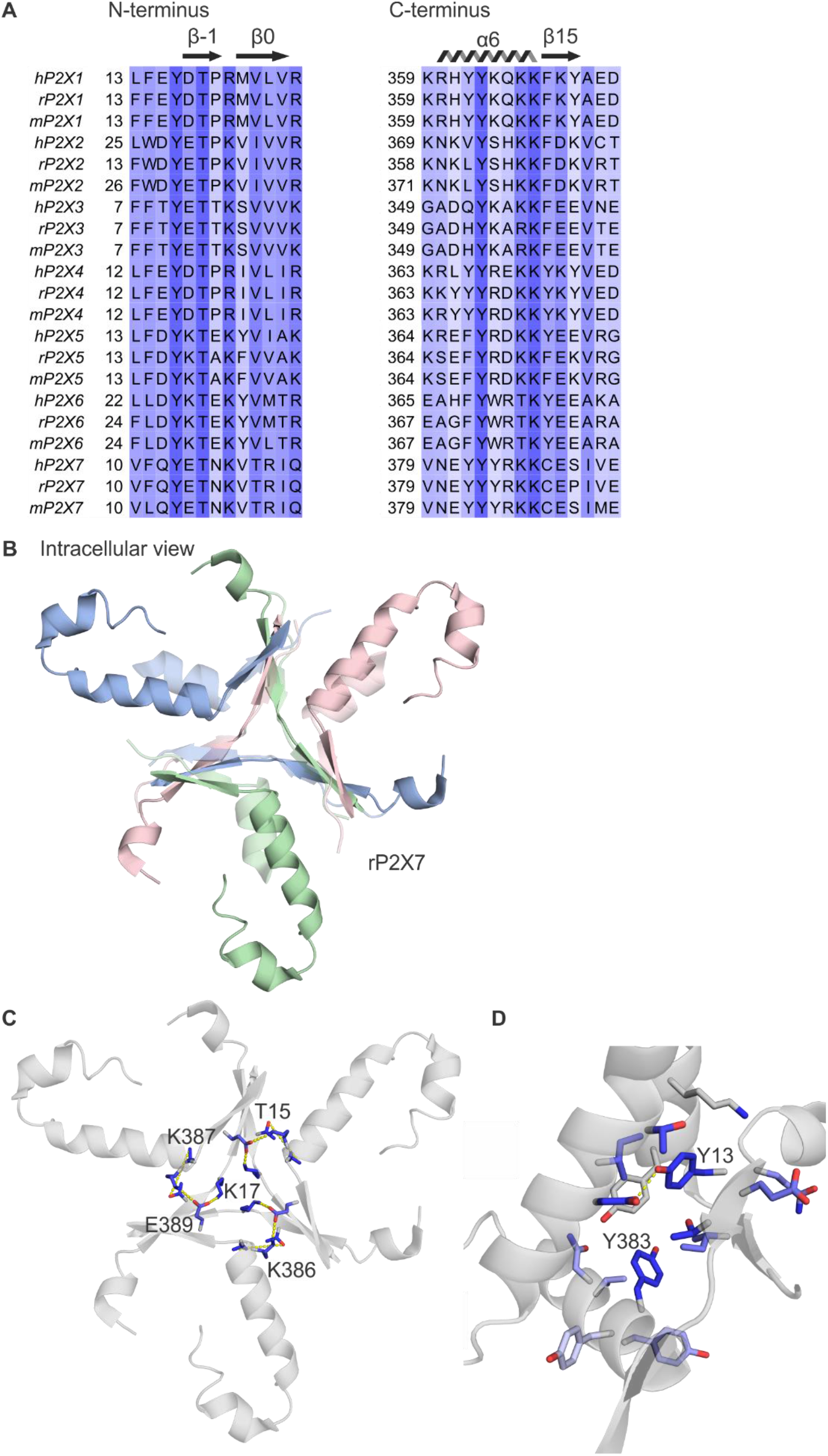
Conservation of the cytoplasmic cap in P2X receptor channels. **(A)** Multiple sequence alignment of all human, rat, and mouse P2X subtypes, truncated to show residues that contribute to the cytoplasmic cap in available structures of human P2X3 and rat P2X7. Columns are colored by the alignment quality score calculated in Jalview based on BLOSUM 62 scores. **(B)** Intracellular view of cytoplasmic cap from the structure of rP2X7 in complex with ATP (PDB ID: 6u9w) in the same orientation as Fig. 2 panel A, with the hP2X3Slow cap overlaid for comparison. **(C)** Intracellular view of cytoplasmic cap from the structure of rP2X7 in complex with ATP in the same orientation as panel B, showing intersubunit side chain interactions between K386 and E389 (3.1-3.3 Å apart); T15 and K387 (7.2-7.4 Å apart, though other rotamers would bring them close enough for hydrogen bonding similar to T12 and K357 in P2X3_slow_); and K17 and E389 (2.4-2.6 Å apart) in yellow. Side chains are colored according to alignment quality score calculated from the multiple sequence alignment in panel A, where highly conserved residues are colored in blue and poorly conserved residues are colored in white. Alignment quality score calculated in Jalview based on BLOSUM 62 scores. **(D)** View from the lateral fenestration of rP2X7 of two conserved tyrosine residues (Y13 and Y383) and the side chains of residues within 4.0 Å. Y13 and D352 of the neighboring subunit are 2.9-3.0 Å apart. This view is from the same orientation as Fig. 2H.

**Figure 2 – Figure Supplement 2.**
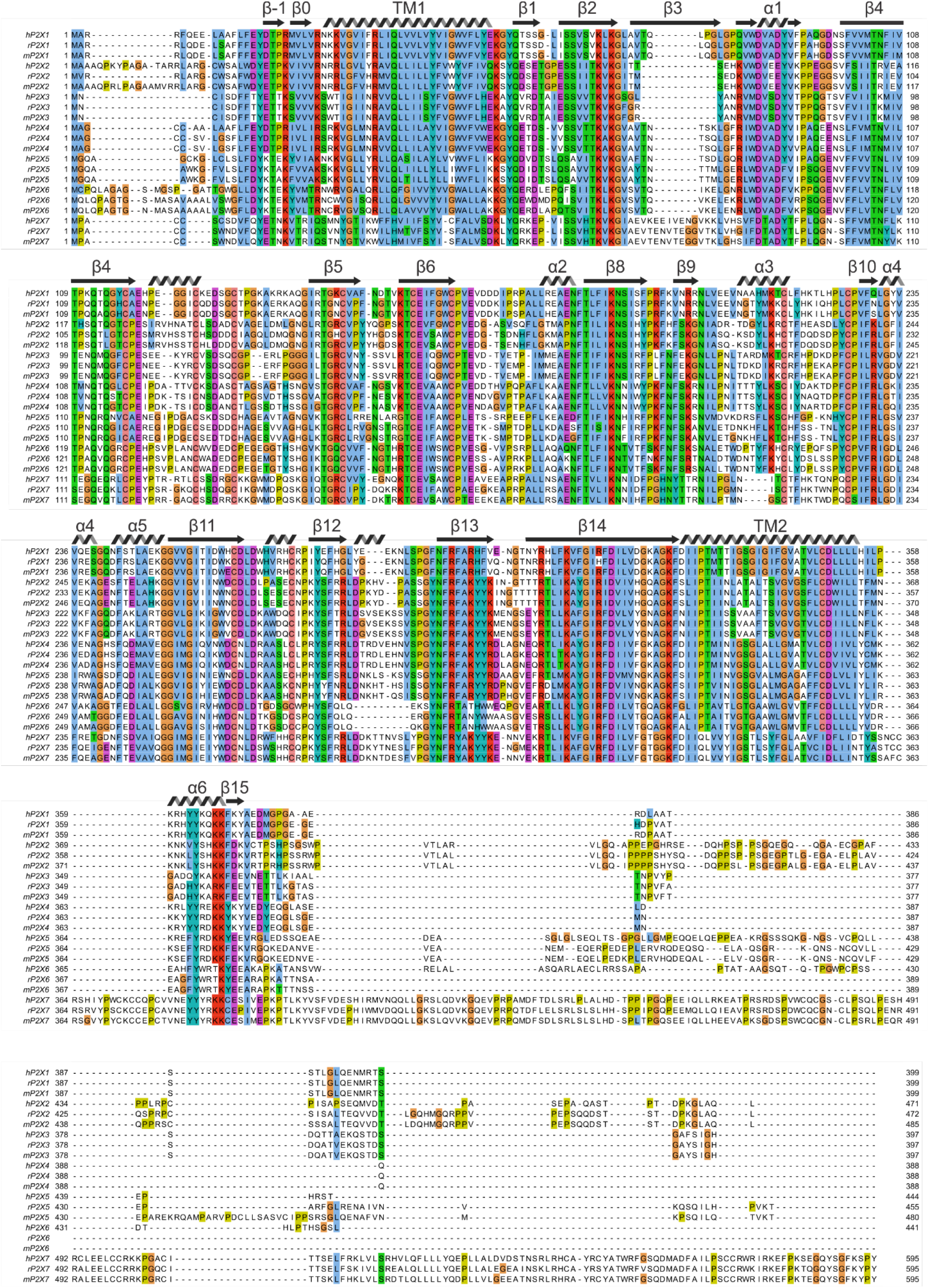
Multiple sequence alignment of human, rat and mouse P2X receptor channels. Multiple sequence alignment obtained using TCoffeeWS via Jalview with residues are colored using Jalview’s ClustalX color scheme, which is based on side chain character and conservation at that position of the alignment. Secondary structural elements are from hP2X3_slow_. Uniprot accession numbers are as follows: hP2X1, P51575; rP2X1, P47824; mP2X1, P51576; hP2X2, Q9UBL9; rP2X2, P49653; mP2X2, Q8K3P1; hP2X3, P56373; rP2X3, P49654; mP2X3, Q3UR32; hP2X4, Q99571; mP2X4, Q9JJX6; hP2X5, Q93086; rP2X5, P51578; mP2X5, B7ZND5; hP2X6, O15547; rP2X6, P51579; mP2X6, O54803; hP2X7, Q99572; rP2X7, Q64663; mP2X7, Q9Z1M0.

**Figure 3 – Figure Supplement 1.**
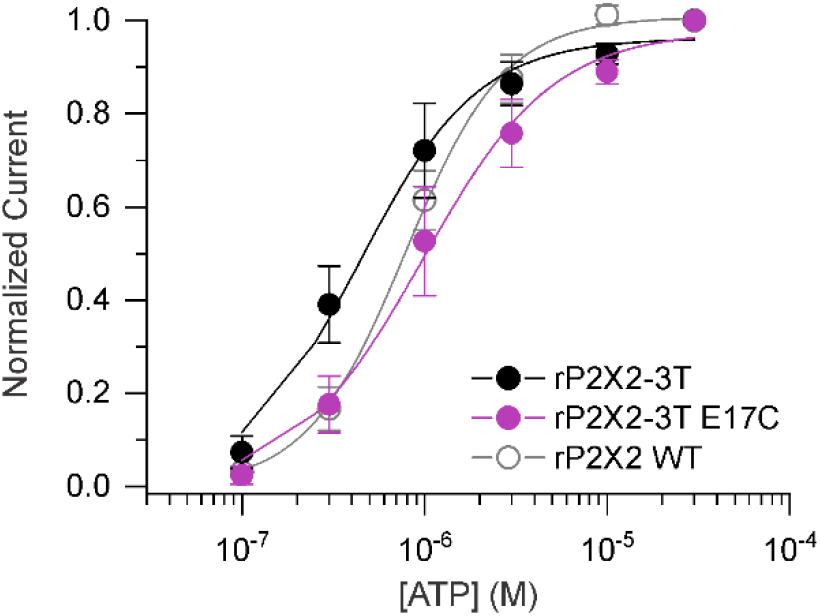
Concentration-dependence for activation of rP2X2 constructs. Normalized concentration-dependence for ATP activation of rP2X2-3T (black filled circles, n=3), E17C rP2X2-3T (purple filled circles, n=3) and WT rP2X2 (gray open circles, n=4) in solutions without divalent ions. Smooth curves are fits of the Hill equation to the data with EC_50_ and n_H_ values of 0.44 ± 0.6 μM and 1.3 ± 0.2 for rP2X2-3T, 1.0 ± 0.1 μM and 1.2 ± 0.2 for E17C rP2X2-3T and 0.8 ± 0.1 μM and 1.7 ± 0.1 for WT rP2X2. Error bars are SEM.

